# Neurobehavioral Effects of “Dry Hit” Nicotine E-Cigarette Vapor Inhalation in Adolescent Wistar Rats

**DOI:** 10.64898/2026.03.26.714509

**Authors:** Aliyah Ogden, Sophia Wright, Sri V. Kasaram, Samuel Moutos, Chris Wernette, Mariana I.H. Dejeux, Benjamin A. Schwartz, Christie M. Sayes, Jacques D. Nguyen

## Abstract

“Dry Hitting” is a unique phenomenon of e-cigarette use that has been shown to produce toxic chemical degradants and byproducts. Although it is widely understood that nicotine exposure during adolescence impacts neurobiological and behavioral function, little is known about how dry hitting may impact users. We hypothesized that subjects repeatedly exposed to nicotine dry hit vapor would exhibit distinct behavioral responses compared with saturated nicotine vapor and would differentially alter the expression of perineuronal nets (PNNs) in the rodent brain. Using a customized system of e-cigarette vapor inhalation, adolescent male Wistar rats (PND 31-40) received vaporized nicotine (30 or 60 mg/mL; ∼2.5-3 mL/cage), nicotine with dry hits (60 mg/mL; 1.75-2 mL/cage), or propylene glycol (PG) vehicle for 30 minutes over 7 daily sessions. Locomotor activity, antinociception, and elevated plus maze testing were used to assess behavioral response to drug intoxication and tolerance. Immunohistochemistry was used to identify *Wisteria Floribunda Agglutinin* (WFA)-positive PNN structures in the amygdala and insular cortex. Rats exposed to dry hits exhibited behavioral responses (locomotor sensitization, antinociception) similar to those of rats exposed to saturated nicotine vapor, but spent more time in the open arms of the elevated plus maze. Immunohistochemical analyses confirmed significantly greater WFA intensity in the central nucleus of the amygdala, but not the basolateral amygdala or insular cortex, of rats exposed to dry hits. Overall, these data confirm the impact of dry hit vapor on behavioral responses and perineuronal net expression in rats during adolescence.

## 1. Introduction

Nicotine exposure during early life contributes to future use and addiction liability during adulthood. Although cigarette smoking has decreased in recent years, e-cigarettes remain popular, especially amongst the youth (Marynak, 2025; Meza et al., 2020). E-cigarettes are also the most common form of nicotine use in teens, with nearly 6% of middle and high school students using e-cigarettes (Cardenas et al., 2021; Park-Lee et al., 2024). The side effects of e-cigarette device use on health are relatively unknown, particularly because of the variety of chemicals found in the devices and the different patterns of regular use among the youth (Hair et al., 2023; Park et al., 2022). Importantly, repeated or chronic use of addictive drugs such as nicotine can result in both neural and behavioral adaptations, including changes in neurotransmitter signaling, changes in glial cell function, and behavioral tolerance (Kulbe et al., 2023; Thibeault et al., 2025). Studies have found that the nicotine content in e-cigarettes (e.g., JUULs) can be similar to that of 18 cigarettes, and individuals using e-cigarettes have a much higher nicotine dependence than traditional tobacco smokers (Jankowski et al., 2019; Prochaska et al., 2022). Thus, preclinical models of vapor administration are critical to understanding how e-cigarettes modulate neurobehavioral outcomes (Nguyen et al., 2016).

“Dry hitting” is a phenomenon unique to e-cigarette use and occurs when the liquid level in a refillable e-cigarette cartridge gets too low, causing the coil inside the tank to overheat and burn (Beard et al., 2024). Whereas e-liquid components can change as a result of thermal decomposition and reactions (Dada et al., 2022), the e-liquid vapor following dry hits and aged coils can contain toxins, including ketene, formaldehyde, acetylaldehyde, acrolein, or acetone (Beard et al., 2024; Farsalinos et al., 2015; Goto et al., 2022). When co-administered with nicotine, acetaldehyde has been shown to enhance the behavioral effects of nicotine in adolescent subjects (Cao et al., 2007), including nicotine self-administration in rats (Belluzzi et al., 2005). Moderate doses of acetaldehyde administered intravenously have been shown to increase locomotor activity and rearing episodes in adult rats (Correa et al., 2003). The high temperatures associated with the production of harmful levels of ketene, which can cause severe lung damage and lead to the development of E-cigarette or Vaping Product Use-Associated Lung Injury (EVALI) (Narimani & da Silva, 2020). Furthermore, exposure to dry-hit vapor decreases cellular metabolism in alveolar epithelial cells compared with saturated conditions without dry hits (Beard et al., 2024).

Perineuronal nets (PNNs) are specialized netlike structures composed of extracellular matrix (ECM) aggregates and glycoproteins, which typically surround the cell bodies and dendrites of GABAergic, parvalbumin (PV) -containing inhibitory interneurons and can be modulated by exposure to nicotine (Bosiacki et al., 2019; Duncan et al., 2019; Honeycutt et al., 2024; Slaker et al., 2016; Vazquez-Sanroman et al., 2017). The formation of PNNs is associated with the maturation of the cortex, as they often form during the critical period of development, and is usually experience-dependent (Bosiacki et al., 2019; Brown & Sorg, 2023; Duncan et al., 2019; Honeycutt et al., 2024; Peeters & Grueter, 2024). These structures are responsible for limiting synaptic plasticity in adulthood, maintaining existing synaptic connections, and protecting neurons from oxidative stress (Bosiacki et al., 2019; Brown & Sorg, 2023; Duncan et al., 2019; Honeycutt et al., 2024). PNNs are necessary for creating and maintaining several types of memory, including drug-related memories (Bosiacki et al., 2019; Duncan et al., 2019; Guarque-Chabrera et al., 2022; Slaker et al., 2016). Drug-related memories include drug-related cues and drug-related context memories, which are strengthened over time with repeated use of the drug. Drug exposure is known to alter PNN expression; for instance, nicotine exposure resulted in more intense expression in the insula (Guarque-Chabrera et al., 2022; Honeycutt et al., 2024; Slaker et al., 2016). PNNs are a novel focus in addiction neuroscience and are incredibly important in neuroplastic mechanisms.

The amygdala and the insular cortex are involved in nicotine’s effects on motivation, reward, and reinforcement, potentially through dynamic regulation of perineuronal nets (PNNs) (Honeycutt et al., 2024; Slaker et al., 2016). PNNs within the amygdala are associated with fear and addiction-related behaviors and show low expression in rodents but high expression in humans (Slaker et al., 2016). PNNs in the insula are considered to contribute to maladaptive drug-associated behaviors (Tomek et al., 2020). Repeated activation of α7 nAChRs during the development of PNNs may also regulate expression of PV-positive neurons (Tomek et al., 2020). Overall, the objective of this study was to determine if e-cigarette nicotine or dry hit exposure during adolescence results in changes in expression of PNNs, which may regulate behavioral responding and underlie vulnerability to compulsive drug use during adolescence.

## 2. Methods

### 2.1 Subjects

The subjects for the study were male Wistar rats (N=80), aged approximately 4-5 weeks old (Charles River; Wilmington, MA). All animals were housed and handled in accordance with the Baylor University Institutional Animal Care and Use Committee. Animals were provided ad libitum access to food and water and were housed on a reverse 12:12 light cycle. Behavioral experiments took place during the scotophase under red light.

### 2.2 Drugs

Nicotine ditartrate dihydrate (Fisher Scientific Company; Bridgewater, NJ, USA) was dissolved in propylene glycol (PG) (Sigma-Aldrich Corporation; St. Louis, MO, USA) and administered via passive vapor inhalation once daily for seven consecutive days. The subjects were exposed to various concentrations of e-liquid solution, including PG, nicotine (30 and 60 mg/ml), and nicotine (60mg/ml) with a dry hit. A separate control group that received only air showed no significant difference from the PG control group on behavioral assessments (**Supplemental Materials**).

### 2.3 Passive Vapor and Dry Hit Procedure

A vapor delivery system and e-vape controller (La Jolla Alcohol Research, Inc.) was used for passive nicotine vapor delivery and was comprised of a custom sealed plexiglass chamber (38.25 in x 23 in x 10.5 in), which accommodated four cages (1 rat/cage), connected to an atomizer (Geekvape Z Top Airflow Sub-ohm Tank; Geekvape Z series coil, 0.4-ohm, 50∼60W). An external wall vacuum system was used to exhaust vapor at a rate of 4-5 L per minute. Two barometers were used to monitor pressure differences between the inside of the plexiglass chamber and the outside environment. Vapor was delivered in once daily sessions for 30 minutes using five 10-second puffs every five minutes (Gutierrez et al., 2021; Gutierrez, Nguyen, et al., 2024; Gutierrez et al., 2022; Kulbe et al., 2023; Nguyen et al., 2016; Nguyen, Creehan, Grant, et al., 2020; Nguyen, Creehan, Kerr, et al., 2020; Nguyen et al., 2021).

Throughout the session, the e-liquid tank was consistently refilled during the five-minute break periods to ensure no dry hits occurred. After the 30-minute session, there was a 4-minute clear-out period during which the vacuum was turned on to ensure that vapor from the chamber was exhausted. The “Dry Hit” condition was defined as when the coil inside of the vapor tank produced a bright orange glow (**Figure 1**). A new coil was used at the start of each dry hit session, and cages were changed between groups. This once-daily passive delivery consisted of 30-minute sessions with sets of 10 2-second vapor puffs, delivered every 2 minutes (Beard et al., 2024). The vapor delivery time in this paradigm is equivalent to that previously described in the saturated passive vape procedure. At the start of the session, the tank was filled with 1mL of drug solution. Animals were given dry hits of a 60mg/ml nicotine solution. After each consecutive set of ten vapor hits, the tank was filled with 0.5mL of solution. This ensures the coil will not fully burn out when no liquid is present, but there will also not be too much liquid, which would prevent a dry hit. Coils were reused between sessions of the same drug unless there was a mechanical or functional reason, they needed to be changed. Different coils and tanks were used for each condition, and the tanks were cleaned thoroughly at the beginning and end of each experimental day. Cages were changed between groups.

**Figure 1.**
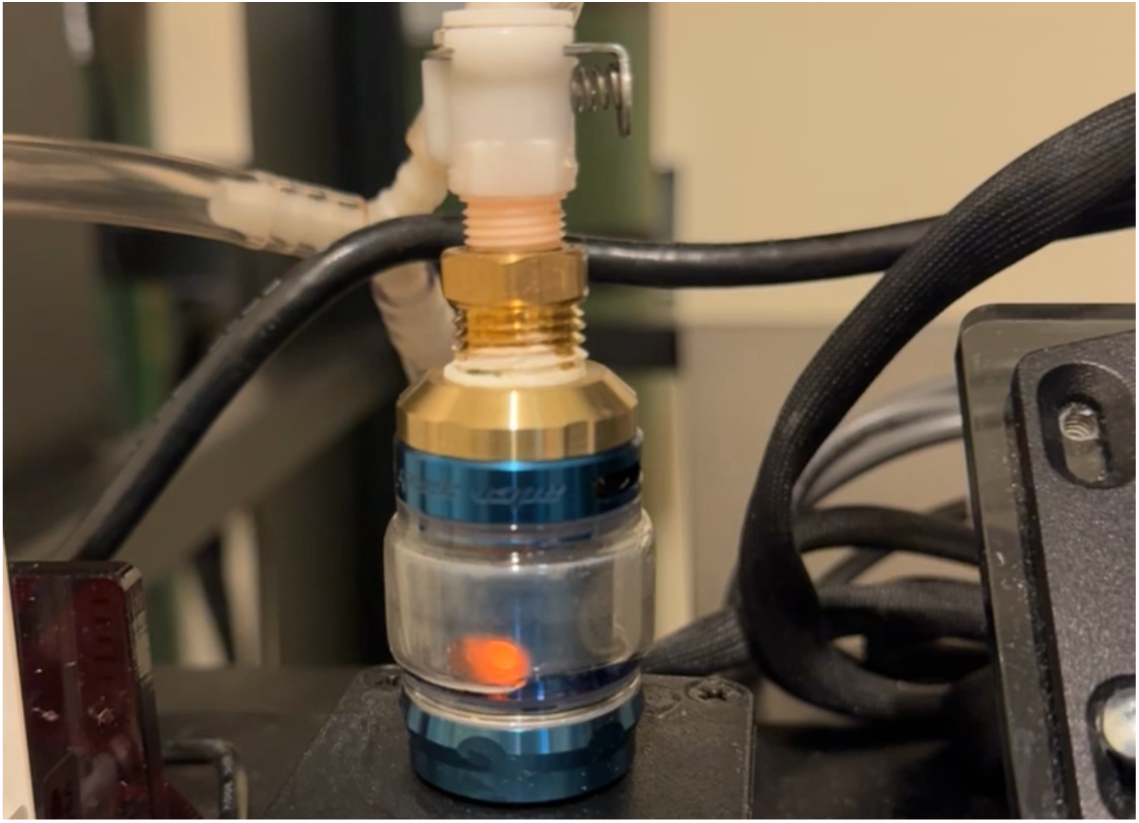
E-cigarette tank during a “dry hit” episode. When activated, the unsaturated coil glows a bright orange due to overheating.

### 2.4 Behavioral Procedures

#### 2.4.1 Antinociception Test

The tail-withdrawal assay was used to evaluate the antinociceptive effects of nicotine and dry hit vapor. Using a VWR® Digital General Purpose Water Bath, one inch of the rat’s tail was submerged in water at 54(±1)°C, and two different timers recorded how long the tail was submerged, with a cut-off time of 15 seconds. The average of the two times was used in the analysis. The longer the tail is in the water, referred to as latency, the greater the drug’s antinociceptive effect. This assay was performed on the first and final days of drug treatments. Tail withdrawal was performed immediately before and 5 minutes after vapor administration. The measurement before the drug exposure was used as a baseline. It can be inferred that nociceptive tolerance developed if there was a significant decrease in latency after drug exposure on the first and last days of vapor exposure.

#### 2.4.2 Locomotor Activity Test

A locomotor activity (LMA) test was used to measure the effects of nicotine and dry hit vapor. This test occurred 15-20 minutes post-vapor administration and lasted 60 minutes. The locomotor boxes were environment-controlled chambers (SuperFlex Open Field System Environmental Control Chambers; OmniTech Electronics, Inc.). The lights remained off for the entirety of the hour in which locomotor behavior was measured. The data that were collected included total distance traveled and ambulatory activity count (Fusion Software v6.5). Stereotypic episode count, vertical episode count, rest episode count, stereotypy time, and rest time were also quantified (Supplemental Materials). Ambulatory activity count was defined as the number of beam breaks while the animal is moving and excludes stereotypic behaviors. Total distance was defined as the total distance in centimeters the animal traveled during the session. Stereotypic episodes were defined as distinct periods of time, separated by at least 1 second, during which the animal exhibited stereotypic behaviors. Vertical episodes were defined as when the animal breaks a vertical beam and are separated by at least 1 second of the animal below the vertical beam. Rest episodes occurred when the animal was inactive for more than 1 second. Stereotypic time was the amount of time during which stereotypic behavior was observed. Rest time was the amount of time during which inactivity was observed.

#### 2.4.3 Elevated Plus Maze

The elevated plus maze (EPM) was used to identify anxiety-related behaviors due to acute nicotine and dry hit vape exposure in a subset of rats (N=8-12). This test was completed immediately after tail withdrawal and immediately before LMA. A four-armed maze located 50 centimeters off the ground, consisting of two open and two closed arms located at right angles from one another, was used for this test. The animal was placed in the center of the maze and was allowed to explore for 5 minutes. This data was recorded and then analyzed using EthoVision XT Software v14. Recordings were 5 minutes ±10 seconds. The analyzed data included the amount of time spent in each of the open arms, closed arms, and the center of the maze. Due to slight variability in video length, values were standardized to percentages prior to data analysis. All data are reported as a percentage of the total time spent in the maze in each of the three areas.

### 2.5 Tissue Collection for Immunohistochemistry

24 hours after the final session of drug exposure, isoflurane was used to anesthetize subjects before transcardial perfusion with 4% paraformaldehyde (PFA), followed by rapid decapitation. The collected brains were postfixed in 4% PFA for 24 hours, then transferred to a sucrose solution until they sank. The tissue was then flash-frozen in isobutane cooled with dry ice before being stored in the -80 ^°^C freezer prior to analysis.

### 2.6 Immunohistochemistry

Tissue from the insular cortex and amygdala was collected as coronal sections (30 μm thickness) using a CryoStar NX50 (Epredia), and areas of interest (basolateral and central amygdala, -2.04mm to -2.52mm from bregma; insular cortex, 2.52mm to 2.16mm from bregma) were determined using (Paxinos & Watson, 2014). Slices were stored in a 0.01% sodium azide solution until immunohistochemistry (IHC) analysis of PNN and PV+ interneuron expression. 0.3% Triton-X, 2% goat serum, 1% BSA was used as a blocking agent; tissue slices were incubated in primary antibodies Wisteria Floribunda Lectin Antibody, Fluorescein (FL-1351-2; Vector; 1:500), Anti-NeuN Antibody, clone A60MAB377 (Sigma-Aldrich; 1:1000), and Parvalbumin Polyclonal Antibody, Unconjugated, Host: Rabbit / IgG (PA1-933; Invitrogen; 1:10,000) overnight. The second day of IHC involved incubating the slices in the secondary antibodies (NeuN: Goat anti-mouse IgG (H+L) Cross-Adsorbed Secondary Antibody, Alexa Fluor 568, 1:200; PV: Goat anti-Rabbit IgG (H+L) Cross-Adsorbed Secondary Antibody, Alexa Fluor 647, 1:1000) and then mounting the slices on microscope slides (FisherScientific). The following day, the slides were cover-slipped with polyvinyl alcohol mounting medium (Sigma-Aldrich) and left to dry for 24 hours before imaging. WFA expression and PV expression (**Supplemental Materials**) were quantified for this study, with three subjects excluded due to experimental exigencies. All samples were imaged using a Keyence BZ-X1000 Fluorescence Microscope at a 20x objective. Images were then uploaded and analyzed using Polygon AI software (ImageJ v1.54r; NIH, Bethesda, MD). WFA^+^ and PV^+^ cells were counted using a pre-existing WFA- and PV-specific detection model and manually corrected when false negatives or positives were identified and corrected by a blinded researcher. Mean (±SEM) WFA^+^ and PV^+^ density were quantified based on cell count divided by the area for each subregion between two hemispheres of each individual subject and represented as (count/mm^2^). Mean (±SEM) WFA^+^ and PV^+^ Intensity were quantified based on individual cell intensity values per animal within the outlined area for each subregion and represented as arbitrary units (AU). Background fluorescence was subtracted from each image.

### 2.7 Statistical Analyses

Data were analyzed using analysis of variance (ANOVA) to identify significant main effects of Treatment, Time (comparing day 1 and day 7, changes over time in the locomotor analysis session, and before and after drug exposure), and the Treatment × Time interaction. Results were considered significant at a p-value of less than 0.05. If a significant main effect was identified, a Tukey’s post post-hoc analysis for multiple comparisons was performed to identify differences between individual conditions.

## 3. Results

### 3.1 Repeated dry hit exposure potentiated tolerance to nicotine vapor-induced changes in spontaneous locomotor activity

Locomotor activity was determined by analyzing the mean (±SEM) total distance traveled and ambulatory activity. These data were collected as ten-minute time bins and as cumulative data per session. Time bin data were analyzed using a two-way ANOVA and a Tukey’s post-hoc multiple comparisons test. The cumulative data were analyzed using a one-way ANOVA and Tukey’s post-hoc multiple comparisons test. **Figure 2A** shows the total distance traveled over the hour-long LMA test, represented as ten-minute time bins, on the first day of passive nicotine vapor exposure. The statistical analyses revealed a significant Time × Drug interaction [F(15,300)=4.380; *p*<0.0001] and a significant Time effect [F(5, 300)=83.94; *p*<0.0001]. The Drug effect was not significant [F(3,60)=1.534; *p*=0.2149]. Post-hoc analyses revealed significant differences in the 10-minute time bin. In the first ten minutes of the test, there were significant differences found between PG Control and 60mg/ml (*p*<0.0001), PG Control and dry hit (*p*<0.0001), 30mg/ml and 60mg/ml (*p*=0.0016), and 30mg/ml and dry hit (*p*=0.0266). In the first 10 minutes, passive nicotine vapor inhalation dose-dependently repressed locomotor activity. All treatment groups showed a significant decrease in total distance traveled per time bin after the first 10 minutes. **Figure 2B** shows the sum of all time bins for each condition on Day 1, which represents the total distance traveled over the hour. ANOVA and post-hoc analysis revealed no significant differences [F(3,60)=1.534; *p*=0.2149].

**Figure 2.**
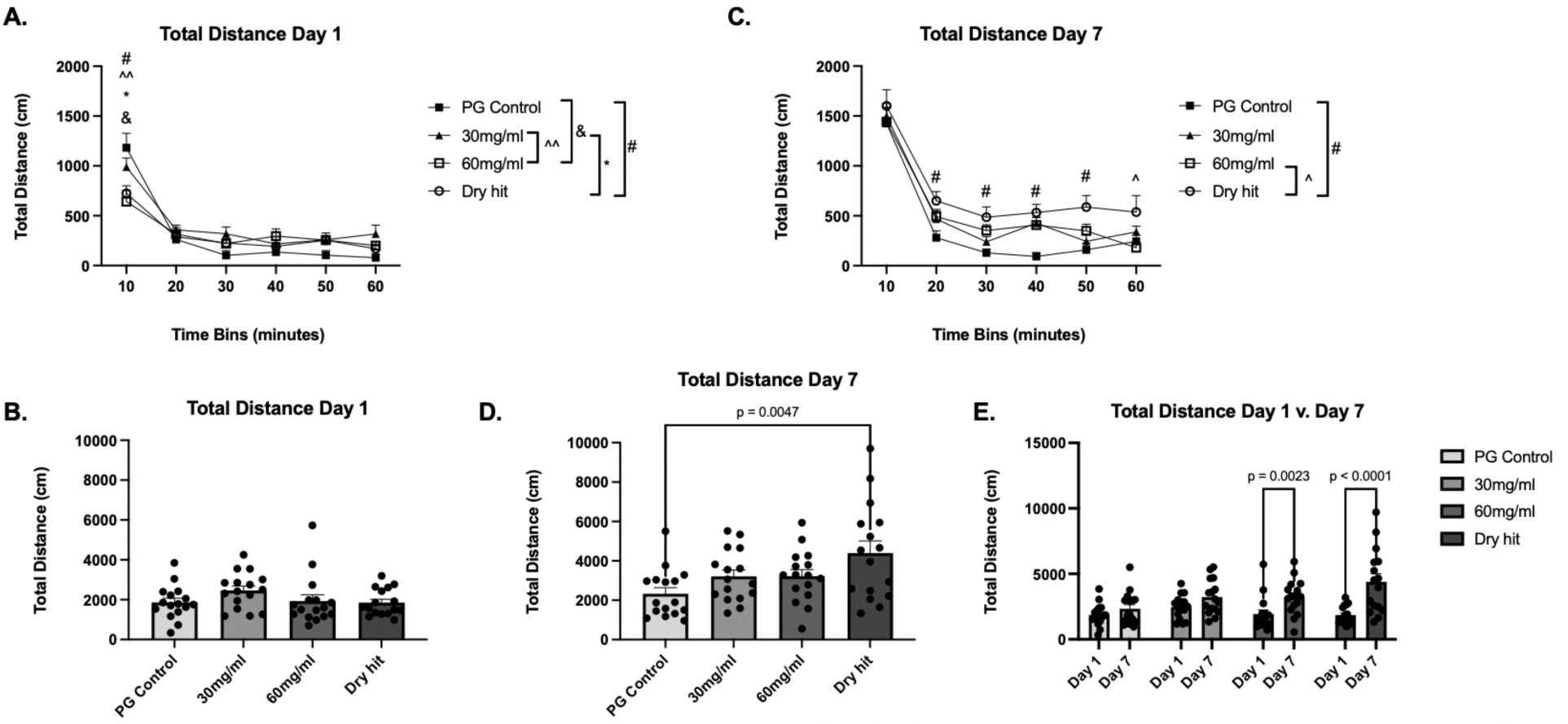
Mean (N=16 per group; ±SEM) total distance traveled during a 1-hour session. (A, C) Total distance traveled on Day 1 or 7 of repeated vapor exposures in ten-minute time bins for the hour-long locomotor assessment period. (B, D) Sum of all time bins over on Day 1 or 7. (E) Comparison of total distance traveled on Day 1 vs. Day 7 across treatment conditions. Significant differences are indicated by *(dry hit and 30mg/ml), ^(dry hit and 60mg/ml), ^^(30 and 60 mg/ml), #(dry hit and PG), or &(60mg/ml and PG).

As shown in **Figure 2C**, the total distance during the hour-long testing session on Day 7 of repeated nicotine vapor exposure, when separated into 10-minute time bins, showed a robust change in activity across conditions. A 2-way ANOVA of the data revealed a significant Time effect [F(5,300)=179.1; *p*<0.0001] and a significant Drug effect [F(3,60)=4.144; *p*=0.0098]. The interaction of Time × Drug was not significant [F(15,300)=1.140; *p*=0.3195]. A Tukey’s post-hoc multiple comparisons test revealed significant differences within time bins and across treatment groups. There were differences between PG control and dry hit in the 20-minute time bin (*p*=0.0294), 30-minute time bin (*p*=0.0385), 40-minute time bin (*p*=0.0057), and 50-minute time bin (*p*=0.0077). There was a significant difference between the 60mg/ml and the dry hit groups in the 60-minute time bin (*p*=0.0376).

**T**he sum of all time bins for each condition is shown in **Figure 2D**; the data represent the total distance traveled over the hour-long session on Day 7 of repeated nicotine vapor exposure. The one-way ANOVA analysis revealed a significant Time × Drug interaction [F(3, 60) = 4.144, *p* < 0.0098]. Post-hoc analyses showed significant differences between PG control and dry hit (*p*=0.0047). Animals in the Dry Hit group traveled a significantly greater total distance during the hour-long session on day 7 than any other treatment group, including those receiving an equivalent nicotine dose. The total distance traveled during the hour-long session on Days 1 and 7 of drug exposure for each treatment group is compared in **Figure 2E**. A two-way ANOVA revealed a significant interaction of Time × Drug [F(3,60=5.118; *p*=0.0032], a Time effect [F(1,60)=38.68; *p*<0.0001]. There was no significant Drug effect [F(3,60)=2.601; *p*=0.0603]. Post-hoc analyses showed no significance between groups on Day 1. However, on Day 7, there were significant differences between PG control and dry hit (*p*=0.0002). There was no difference between PG control Day 1 and Day 7 (*p*=0.2575) and 30mg/ml Day 1 and Day 7 (*p*=0.0693). Significant differences were found between 60 mg/ml Day 1 and Day 7 (*p*=0.0023), and dry hit Day 1 v. Day 7 (*p*<0.0001). These data demonstrate that repeated exposure to nicotine vapor at the doses studied will result in tolerance to the locomotor-suppressing effects observed on Day 1. Dry hits alongside nicotine treatment will enhance this tolerance development effect.

Ambulatory activity data were analyzed in the same manner as the total distance traveled. Day 1 of the assessment, divided into 10-minute time bins, is shown in **Figure 3A**. This data was analyzed using a two-way ANOVA. There was a significant interaction between Drug × Time [F(15,300)=4.421; *p*<0.0001], and a Time effect [F(5,300)=78.42; *p*<0.0001], but there was not a significant Drug effect [F(3,60)=2.401; *p*=0.0765]. Tukey’s multiple comparisons test reveals significant differences in the 10-minute time bin between PG control and 60 mg/ml (*p*<0.0001), PG Control and Dry Hit (*p*<0.0001), 30mg/ml and 60mg/ml (*p*=0.0039), and 30 mg/ml and dry hit (*p*=0.0060). In the 30-minute time bin, PG control and 30 mg/ml differed significantly (*p*=0.0477). A one-way ANOVA was used to analyze the cumulative ambulatory activity, represented in **Figure 3B**. This analysis revealed no significance [F(3,60)=2.401; *p*=0.0765]. Post-hoc analysis also showed no significant differences.

**Figure 3.**
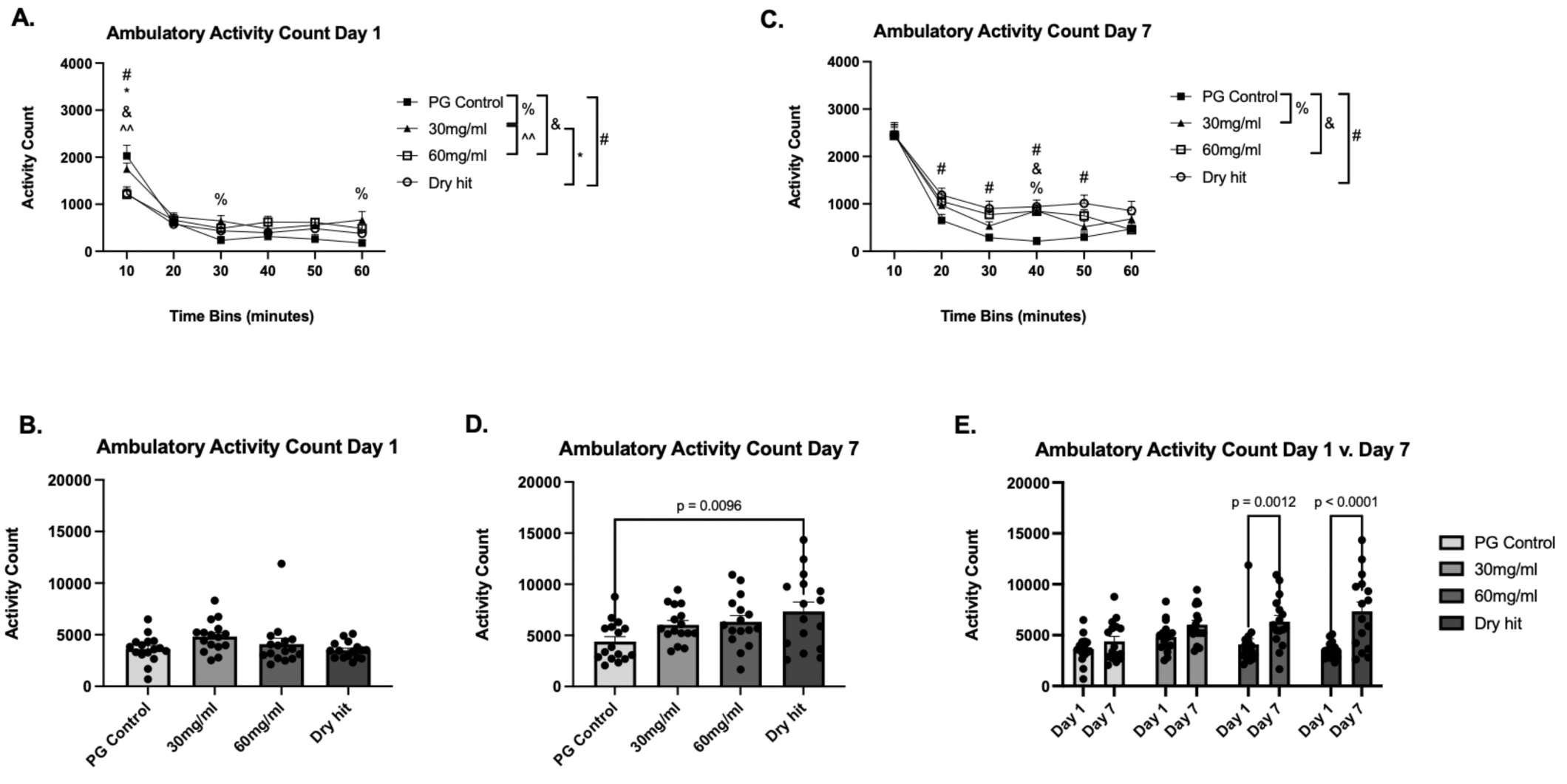
Mean (N=16 per group; ±SEM) ambulatory activity following dry hit or nicotine vapor exposure on Day 1 v. Day 7. (A, C) Ambulatory activity on Day 1 or 7 of repeated vapor exposures in ten-minute time bins for the hour-long locomotor assessment period. (B, D) Sum of all time bins over on Day 1 or 7. (E) Comparison of ambulatory activity on Day 1 vs. Day 7 across treatment conditions. Significant differences are indicated by *(Dry hit and 30mg/ml), ^ (Dry hit and 60mg/ml) ^^(30 and 60 mg/ml), #(Dry hit and PG), % (30mg/ml and PG), or & (60mg/ml and PG).

**Figure 3C** displays Day 7 data collected in 10-minute intervals over an hour-long session, measuring ambulatory activity. A two-way ANOVA confirmed a significant Drug × Time interaction [F(15,300)=2.115; *p*=0.0093], Time effect [F(5,300)=184.1; *p*<0.0001], and Drug effect [F(3,60)=3.658; *p*=0.0173]. Post-hoc analyses for multiple comparisons revealed differences in the 20-minute bin; there were differences between PG Control and Dry Hit (*p*=0.0453). The difference in the 30-minute time bin was between PG control and dry hit (*p*=0.0156). The 40-minute time bin had differences between PG control and 30 mg/ml (*p*=0.0094), PG control and 60 mg/ml (*p*=0.0110), and PG control and dry hit (*p*=0.0022). In the 50-minute bin, a significant difference was found between PG control and dry hit (*p*=0.0029). Lastly, the 60-minute bin displayed no significant differences.

**Figure 3D** shows the cumulative ambulatory activity over the entire hour-long session on Day 7 of repeated nicotine vapor exposure. A one-way ANOVA revealed a significant Drug × Time interaction [F(3,60)=3.658; *p*=0.0173]. Tukey’s test for multiple comparisons revealed a significant difference between PG Control and Dry Hit (*p*=0.0096). Lastly, **Figure 3E** compares the cumulative ambulatory activity counts on Day 1 and Day 7. A two-way ANOVA revealed a significant Drug × Time interaction [F(3,60)=4.374; *p*=0.0075], Time effect [F(1,60)=37.55; *p*<0.0001]. There was no drug effect [F(3,60)=2.657; *p*=0.0564]. Tukey’s multiple comparison test showed no significant differences between groups on Day 1. On Day 7, there was a significant difference between PG control and dry hit (*p*=0.0008). A within-groups comparison between Days 1 and 7 using post-hoc analyses revealed differences in 60mg/ml (*p*=0.0012), and dry hit (*p*<0.0001). The data illustrate a suppression of ambulatory activity on Day 1. Repeated exposure to nicotine vapor resulted in dose-dependent tolerance to that suppression, which was exacerbated slightly by the dry hit condition.

### 3.2 Repeated nicotine vapor inhalation with or without dry hitting produces an acute antinociceptive effect

The tail withdrawal assay was analyzed using a Two-Way ANOVA and tested for multiple comparisons using a Tukey’s Test. All data are represented with the standard error of the mean (SEM). **Figure 4A** shows latency values before and after the first of seven once-daily repeated exposures to nicotine vapor. There was a significant Drug × Time interaction [F(3,28)=4.346; *p*=0.0124] and a significant Time effect [F(1,28)=30.75; *p*<0.0001]. Tukey’s post-hoc test for multiple comparisons revealed there were no significant differences in the pre-test. In the post-test, there were significant differences between PG control and 30 mg/ml (*p*=0.0330), and PG control and 60 mg/ml (*p*=0.0476). Between the pre- and post-tests, there were significant differences in the 30 mg/ml (*p*=0.0005), 60 mg/ml (*p*=0.0003), and dry hit (*p*=0.0024) treatment groups. The data demonstrate that passive nicotine vapor inhalation, with or without a dry hit, induces antinociception in adolescent Wistar rats in a dose-dependent manner. **Figure 4B** illustrates the latency values before and after the final of seven once daily passive nicotine vapor exposures. There was a significant time effect [F(1, 28)=21.50; *p*<0.0001]. A Tukey post-hoc analysis for multiple comparisons revealed significant within-group differences in PG control (*p*=0.0090), 30 mg/ml (*p*=0.0239), and dry hit (*p*=0.0086).

**Figure 4.**
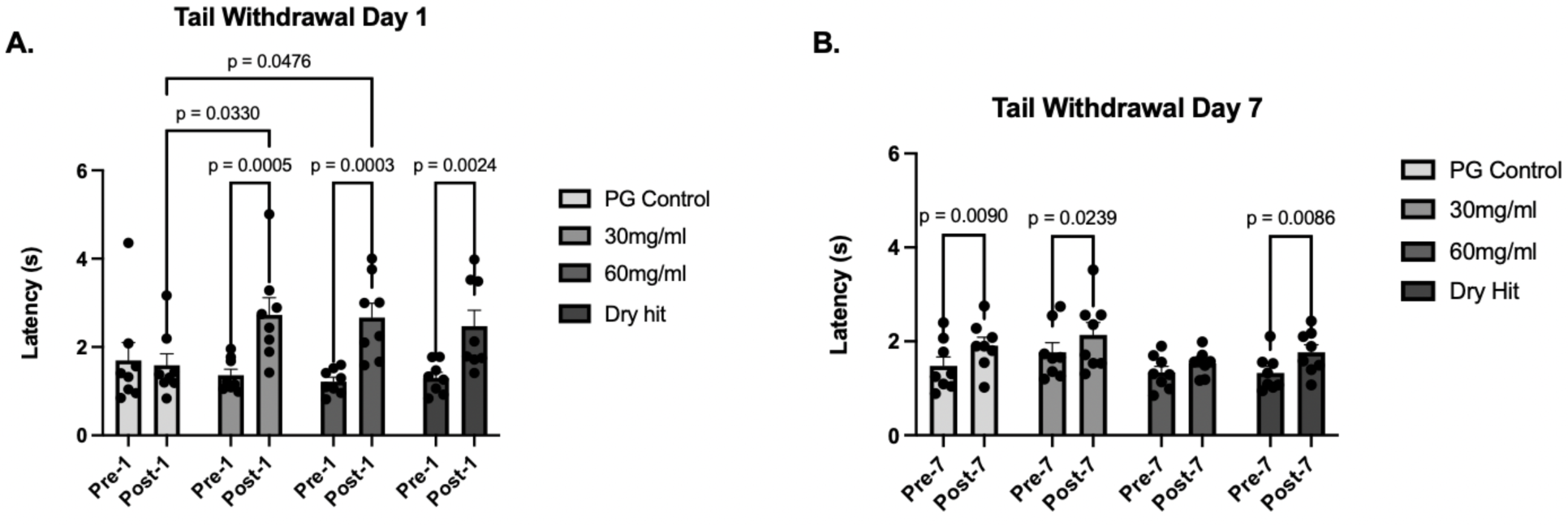
Mean (N=8 per group; ±SEM) tail withdrawal latency pre- and post- nicotine vapor exposure on (A) Day 1 or (B) Day 7.

### 3.2 Repeated dry hit exposure may decrease anxiety-like behavior or increase risky behavior in the elevated plus maze (EPM) test

EPM data were calculated using the percent total cumulative duration in each of the three locations of the maze. **Figure 5A** represents the percentage of time spent in the closed arms, calculated by time in closed arms divided by total time multiplied by 100, on both Day 1 and Day 7. A two-way ANOVA revealed a significant interaction between Time and Drug [F(3,44)=9.783; *p*<0.0001]. Tukey’s Test showed differences between PG Control and Dry Hit on Day 1 (*p*=0.0082). There were significant within-group differences in PG Control (*p*=0.0007), 60mg/ml (*p*=0.0102), and Dry Hit (*p*=0.0141). **Figure 5B** represents the amount of time spent in the open arms, calculated by time in open arms divided by total time multiplied by 100, on Day 1 and Day 7. A two-way ANOVA revealed a significant Time × Drug interaction [F(3, 44) = 9.155, *p* < 0.0001]. Post-hoc analyses revealed a difference between 30mg/ml and 60mg/ml on Day 1 (*p*=0.0428). There were within-group differences in PG control (*p*=0.0003), 30 mg/ml (*p*=0.0062), and dry hit (*p*=0.0374). **Figure 5C** depicts the percentage of time spent in the center of the maze. A two-way ANOVA showed a significant Time × Drug interaction [F(3,44)=4.024; *p*=0.0129]. Tukey’s test revealed a significant difference between PG control and dry hit on Day 1 (p=0.0120). There was a within-group difference between Day 1 and Day 7 in the PG control (*p*=0.0120). The within-group difference of the dry hit group was trending toward significance (*p*=0.0651).

**Figure 5.**
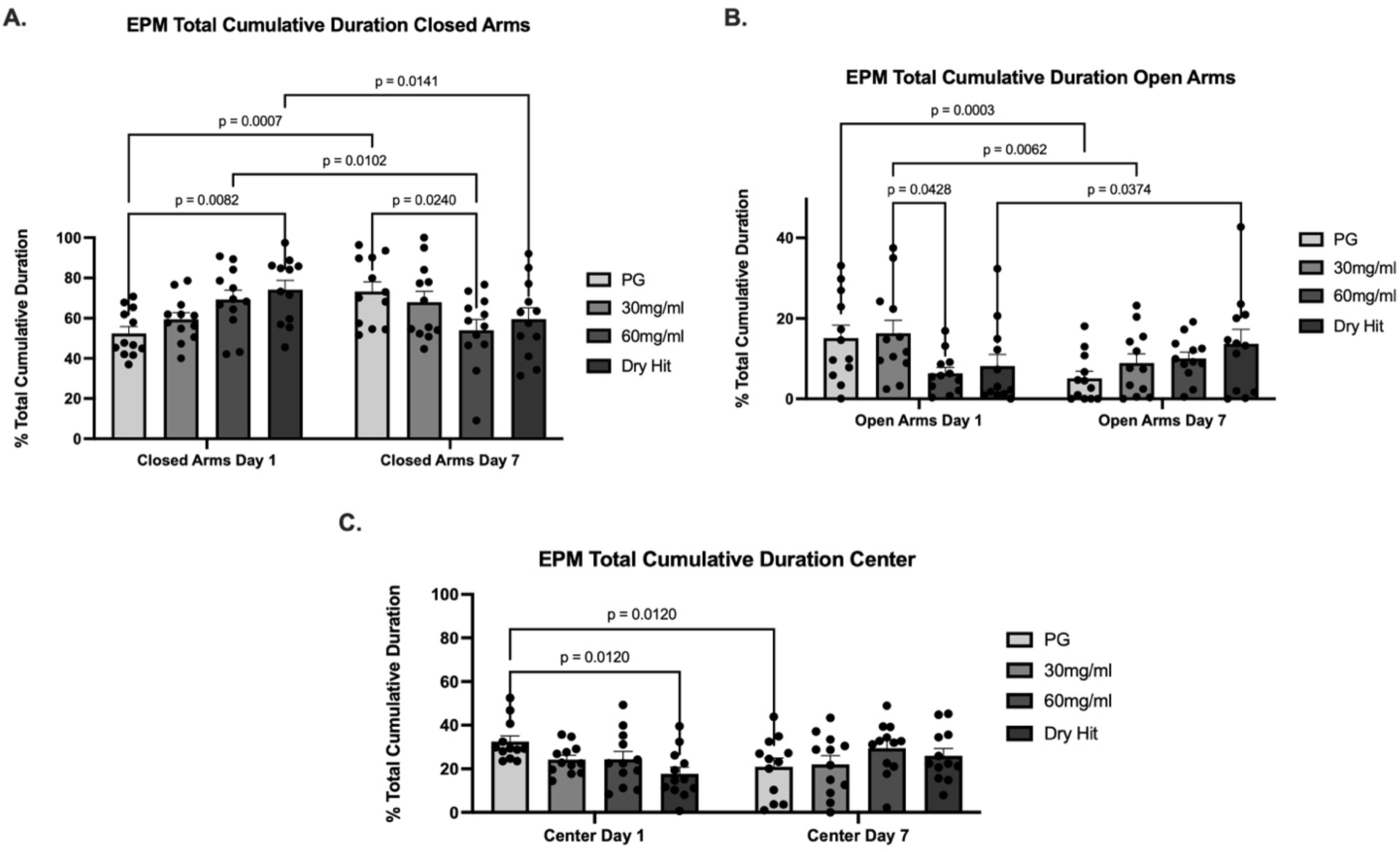
Elevated Plus Maze behavior following repeated dry hit or nicotine vapor exposure during Day 1 and Day 7. Mean (N=12 per group; ±SEM) percent time spent in (A) closed arms, (B) open arms, or (C) center zone on Days 1 or 7.

**Figure 6.**
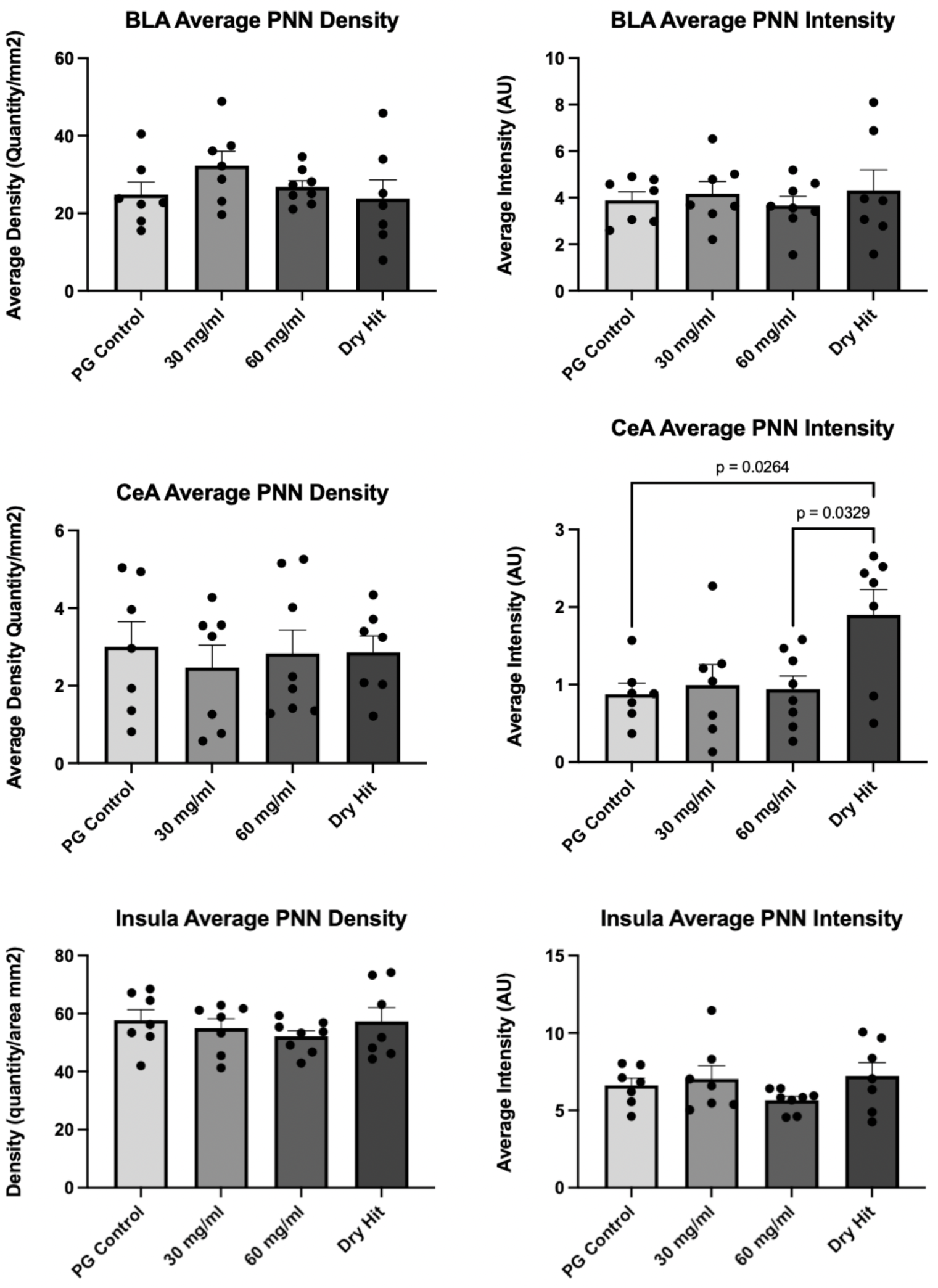
Perineuronal net expression is modulated by dry hit inhalation. Mean (N=7-8 per group; ±SEM) density or intensity of WFA+ tissue in the basolateral amygdala (BLA), central nucleus of the amygdala (CeA) or insular cortex.

### 3.4 Dry hitting increased perineuronal net intensity in the central nucleus of the amygdala (CeA)

Brains that were collected approximately 24-36 hours post vapor-inhalation were analyzed for changes in perineuronal net expression. One-way ANOVA confirmed no significant effect of Drug condition on WFA^+^ density [F(3,25)=1.183; *p*=0.3361] or on WFA^+^ intensity [F(3,24)=0.2672; *p*=0.8484] in the basolateral amygdala (BLA). One-way ANOVA failed to confirm a significant effect of Drug condition on WFA^+^ density [F(3,25)=0.1532; *p*=0.9267]; however, there was a significant effect of Drug condition on WFA^+^ intensity [F(3, 25)=4.172; *p*=0.0159] in the central nucleus of the amygdala (CeA). Post-hoc analyses confirmed a significant increase in intensity following the dry hit condition relative to PG Control (p=0.0264) and to nicotine 60 mg/mL (*p*=0.0329); relative to the 30 mg/ml, there is a trend toward significance (*p*=0.0570). Lastly, one-way ANOVA confirmed that there was no significant main effect of drug condition on WFA^+^ density [F(3,25)=0.5624; *p*=0.6448] or on WFA^+^ intensity [F(3,25)=1.265; *p*=0.3077] in the insular cortex. There were no changes in parvalbumin+ expression in these three regions (data not shown).

## 4. Discussion

Our results indicate that inhalation of dry hits and nicotine e-cigarette vapor produces different neurobehavioral effects in adolescent rats. Whereas repeated nicotine exposure results in neuroadaptations and the development of tolerance (Chellian et al., 2021), with both nociceptive tolerance and locomotor tolerance, repeated dry hit exposure modulated the behavioral effects of repeated nicotine vapor inhalation, including functional tolerance and anxiety-related outcomes (EPM). Nicotine vapor produced a mild antinociceptive effect in a dose-dependent manner at 54^°^C. The animals that received a 60mg/ml dose had a significantly greater antinociceptive effect than those that received a 30mg/ml dose. Nicotine vapor, alongside a dry hit, also has a mild antinociceptive effect, supporting a likely nicotine-induced effect in this condition, and is further supported by a non-significant difference between the dry hit condition and the 30mg/ml or 60mg/ml group. Seven days of repeated passive nicotine vapor exposure resulted in the development of nociceptive tolerance on the final day of testing. This was evidenced by a decrease in drug efficacy for the higher dose of nicotine after the seven repeated vapor exposures and is supported by previous work, which found that seven days of repeated intermittent nicotine exposure in adolescents produces drug dependence (Kallupi et al., 2019).

The lack of a tolerance effect in both the 30 mg/ml and dry hit conditions likely indicates that moderate doses of nicotine or dry hit are required for antinociceptive tolerance, or that dry hit may prevent the development of nociceptive tolerance. As a non-selective nicotinic acetylcholine receptor (nAChR) agonist (Yang et al., 2017), nicotine’s antinociceptive properties may also involve mu opioid receptors activation, which could suggest the effect of dry hits (Barreto de Moura et al., 2019; Carstens et al., 2001; Haghparast et al., 2008; Khalouzadeh et al., 2022; Schochet et al., 2005). Additionally, the α7 nAChR has been implicated in regulating cognitive processes, such as memory and neuronal plasticity, and is highly expressed in the frontal cortex, including the insula (Belluardo et al., 2005; Potasiewicz et al., 2021). The insula is a region in the frontal cortex that modulates behavioral phenotypes induced by drug exposure and integrates sensory, memory, and motivational information (Honeycutt et al., 2024; Loney et al., 2021). Direct nicotine administration to the insula can impact the formation of contextual memories; modulation of the insula by activation of nACHRs on GABAergic interneurons may contribute to the development of substance use disorders (Loney et al., 2021). There were also no changes in WFA+ intensity or density in the insula, which contrasts from previous studies that revealed changes in PNN expression in the insula, due to its role in modulating behavioral phenotypes produced by nicotine exposure, including conditioned place preference, conditioned taste avoidance, and relapse to drug-seeking behaviors after abstinence, and for the integration of sensory and memory (Honeycutt et al., 2024). Adolescent nicotine exposure has also been shown to alter synaptic transmission in the insula (Toyoda & Koga, 2021).

It is expected that nicotine increases spontaneous locomotor behavior during the open field test, an effect seen in both adolescents and adults (Gutierrez, Creehan, et al., 2024; Gutierrez, Nguyen, et al., 2024; Javadi-Paydar et al., 2019); however, the current study reveals acute nicotine vapor at the provided doses results in locomotor suppression, as evidenced by the decreases in ambulatory activity and total distance traveled. This finding contrasts with previous studies investigating the impacts of nicotine vapor on adolescents (Gutierrez, Creehan, et al., 2024; Gutierrez, Nguyen, et al., 2024). Repeated nicotine vapor exposure results in tolerance to locomotor suppression, as indicated by increased total distance traveled and ambulatory activity. This locomotor suppression on day one is enhanced by the presence of a dry hit. Repeated exposure to nicotine vapor at the studied doses will result in tolerance to the locomotor-suppressing effects seen on day one. Dry hits alongside the nicotine treatment will enhance this tolerance development effect. There is a dose-dependent suppression of locomotion, with the development of a locomotor tolerance after repeated exposures. These data are consistent with those collected on ambulatory activity, where there is suppression of ambulation on day one, and a dose-dependent tolerance to that suppression is identified on day seven, which is slightly exacerbated by the addition of a dry hit.

Analysis of EPM data suggests that acute nicotine vapor could potentially increase anxiety-like behaviors and decrease exploratory behaviors. After seven days, however, the amount of time spent in the closed arms decreased in both the 60mg/ml and dry hit groups, suggesting that repeated exposure to nicotine vapor, with or without a dry hit, decreases anxiety-like behaviors. The amount of time spent in the open arms decreased between day one and day seven in the PG vehicle control, suggesting repeated exposure to PG reduces exploratory behavior and increases anxiety-like behavior. The time spent in open arms decreased in the 30mg/ml nicotine vapor group, generally increased in the 60mg/ml group, and significantly increased in the dry hit group, suggesting that repeated exposure to higher doses of nicotine vapor is more likely to result in decreased anxiety-like behaviors and increased exploratory behaviors, while lower doses of repeated nicotine vapor exposure may be anxiogenic. These varying differential effects of nicotine on anxiety-like behavior are in alignment with previous findings represented in the literature (Elliott et al., 2004; Holliday & Gould, 2016).

Lastly, immunohistochemical analyses confirmed that repeated exposure to dry hit vapor significantly increased WFA+ intensity in the central nucleus of the amygdala but not the basolateral amygdala (p<0.05). In support of these findings, early-life stress is known to modulate the density and intensity of perineuronal nets in the amygdala, as well as the morphology of amygdala neurons (Guadagno et al., 2021; Rahimian et al., 2024). PNNs in the amygdala are critical for stabilizing and modulating synaptic activity, particularly in PV-expressing inhibitory interneurons, and are essential for the development of context-dependent memories (Guadagno et al., 2021). Furthermore, CeA PV interneurons are implicated in affective states and anxiety-like behaviors, especially after drug exposure (Wang et al., 2016). These interneurons also modulate amygdala excitability (Ryazantseva et al., 2025).

Work will include further elucidating the potential neurobehavioral impact of specific byproducts from dry hitting, such as acetaldehyde; however, we suspect that due to the role of matrix metalloproteinases (MMPs) in regulating perineuronal nets, it is possible that reactive metal species, such as high levels of zinc, a critical element in MMPs, from dry hit exposure may dysregulate these critical endogenous enzymes (Beard et al., 2024; Verma & Hansch, 2007; Xiao et al., 2024). MMPs can degrade PNNs surrounding PV interneurons; if unregulated or in excess, they can lead to abnormal development and behavior by reducing plasticity and inhibitory modulation (Murase et al., 2017; Wen et al., 2018). Previous studies have shown connections between MMPs produced by astrocytes and microglia and the modulation of neuroinflammation and perineuronal nets (Bosiacki et al., 2019; Cheung et al., 2024). Overall, these data confirm that inhalation of dry hit e-cigarette vapor alters perineuronal net expression in a region-specific manner. Dry hitting and saturated nicotine vapor exposure in rats may differentially alter extracellular matrix structures and immediate early genes integral to neuroplasticity-related learning and behavior. This may indicate a key mechanism underlying an important vulnerability of naïve or inexperienced e-cigarette usage. Future studies should investigate the potential effects of dry hitting and the pharmacokinetics of inhaled nicotine.

Ongoing studies will investigate the role of matrix metalloproteinases in the development of compulsive drug use in adulthood, as well as changes to neuroinflammation and plasticity-related markers after nicotine vapor exposure during adolescence. Because nicotine exposure, including vapor, in adolescence is likely to lead to lasting changes in sensitivity to locomotor stimulation, the lasting behavioral consequences of adolescent nicotine vapor exposure in adulthood will be analyzed as an expansion upon current work. (Gutierrez, Creehan, et al., 2024; Gutierrez, Nguyen, et al., 2024; Kallupi et al., 2019). Data show that adolescent exposure to drugs may result in changes to synaptic signaling and neuropeptide alterations that can persist into adulthood (Flores et al., 2023; Toyoda & Koga, 2021). Therefore, investigations into changes in plasticity from adolescence to adulthood could be particularly important. In conclusion, investigating the effects of dry hitting may be critical to understanding how repeated e-cigarette use during adolescence can lead to enhanced vulnerability and liability for volitional exposure to nicotine vapor.

## 5. Acknowledgements

## Funding and Acknowledgements

This work was funded by support from the United States Public Health Service National Institutes of Health Grant DA047413 (J.D.N.). We would also like to acknowledge the Baylor University Office of Engaged Learning, Undergraduate Research and Scholarly Achievement Grant. The National Institutes of Health/NIDA had no direct influence on the design, conduct, analysis or decision to publish the findings.

